# In vitro analysis of RNA polymerase II elongation complex dynamics

**DOI:** 10.1101/510206

**Authors:** Yoo Jin Joo, Scott B. Ficarro, Yujin Chun, Jarrod A. Marto, Stephen Buratowski

## Abstract

RNA polymerase II elongation complexes (ECs) were assembled from nuclear extract on immobilized DNA templates and analyzed by quantitative mass spectrometry. Time course experiments showed that initiation factor TFIIF can remain bound to early ECs, while levels of core elongation factors Spt4-Spt5, Paf1C, Spt6-Spn1, and Elf1 levels remain steady. Importantly, the dynamic phosphorylation patterns of the Rpb1 C-terminal domain (CTD), and the factors that recognize them, change as a function of post-initiation time, rather than distance elongated. Chemical inhibition of Kin28/Cdk7 blocks both Serine 5 and Serine 2 phosphorylation, affects initiation site choice, and inhibits elongation efficiency. EC components dependent on CTD phosphorylation include capping enzyme, Cap Binding Complex, Set2, and the PAF1 complex. By recapitulating many known features of *in vivo* elongation, this system reveals new details that clarify how EC-associated core elongation factors, chromatin regulators, and RNA processing factors change at each step of transcription.

## Introduction

The RNA polymerase II (RNApII) transcription cycle has three major stages: initiation, elongation, and termination. During elongation, RNApII interacts directly or indirectly with multiple elongation regulatory factors, as well as proteins needed for processing nascent RNA, chromatin modification and remodeling, and eventually termination (Buratowski, 2009). Elongation regulation is critical for proper gene expression, yet much remains unclear about the dynamics of elongation complexes (ECs).

Of the multiple approaches used to identify factors that interact with RNApII ECs, protein biochemistry and yeast genetics have been particularly fruitful. Early *in vitro* elongation experiments using purified RNApII led to discovery of TFIIS, a protein that re-activates stalled, backtracked ECs (Nakanishi et al., 1981; Sekimizu et al., 1976). TFIIS, TFIIF, and the “Polymerase Associated Factor” complex (Paf1C) were purified by their affinity for RNApII (Sopta et al., 1985; Wade et al., 1996). Mammalian DSIF (the Spt4-Spt5 complex) and FACT (Spt16-Pob3-Nhp6 complex) were also isolated by their ability to modulate *in vitro* transcription by purified RNApII factors (Orphanides et al., 1998; Wada et al., 1998). Affinity purifications of these elongation factors uncovered other EC-associated factors such as Spt6, Elf1, and Spn1/Isw1 (Krogan et al., 2002; Prather et al., 2005; Zhang et al., 2008).

These elongation factors were often identified first in genetic screens for transcription regulators. For example, the *SPT* genes were isolated as suppressors of particular transposon insertions (Winston et al., 1984), and many elongation factor mutants are sensitive to drugs that reduce NTP levels, such as 6-azauracil or mycophenolic acid. Chromatin immunoprecipitation experiments show crosslinking of these factors with actively transcribed genes *in vivo*, and several distinct patterns are seen (Kim et al., 2004; Krogan et al., 2002; Mayer et al., 2010).

*In vivo*, RNApII ECs must overcome the inhibitory effect of nucleosomes, but also restore chromatin integrity after passing through (Li et al., 2007; Orphanides and Reinberg, 2000). The elongation factors FACT and Spt6 have histone chaperone activity, and mutations in these genes lead to disrupted chromatin structure, aberrant histone modification, and initiation from cryptic internal promoters (Kaplan et al., 2003; Xin et al., 2009; Youdell et al., 2008). Paf1C is required for H2B ubiquitination, and subsequently several co-transcriptional histone methylations (Krogan et al., 2003a; Van Oss et al., 2016; Wood et al., 2003). The mechanistic details of how these factors function are not yet clear, but recent cryo-EM structures show how several bind to RNApII (Ehara et al., 2017; Vos et al., 2018; Xu et al., 2017b).

Another key component in EC function is the C-terminal domain (CTD) of the RNApII largest subunit, Rpb1. The CTD is comprised of multiple repeats of the heptapeptide sequence Tyr1-Ser2-Pro3-Thr4-Ser5-Pro6-Ser7 (Corden, 2013). During transcription, the CTD undergoes a programmed pattern of phosphorylation and dephosphorylation, generating a ‘CTD code’ that creates binding sites for a variety of proteins needed at different stages of transcription (reviewed in Buratowski, 2009; Corden, 2013). Factors known to bind phosphorylated Ser5 (Ser5P) during early elongation include mRNA capping enzyme, the non-polyA termination factor Nrd1, and the Set1 histone methyltransferase complex. In contrast, mRNA termination factors Pcf11 and Rtt103 and the histone methyltransferase Set2 are coupled to downstream CTD phosphorylation at Ser2P. Mass spectrometry of factors co-immunoprecipitated with different CTD phosphorylations identified additional candidate EC proteins (Ebmeier et al., 2017; Harlen et al., 2016). It is therefore important to understand how the CTD code is generated and used to regulate co-transcriptional processes.

Although reconstitution with purified factors has been essential for identifying the minimal set of EC proteins, transcription *in vivo* is coupled to multiple mRNA processing and chromatin modifying factors that make full reconstitution difficult. Here we use yeast nuclear extracts to better approximate *in vivo* conditions. We previously used quantitative proteomics to analyze RNApII pre-initiation complexes (PICs) (Sikorski et al., 2012). We now extend this analysis to RNApII ECs formed on DNA templates *in vitro*. Mass spectrometry identifies a set of core elongation factors (Spt4-Spt5, Spt6-Spn1, Elf1, and Paf1C), as well as EC-associated histone modifying and mRNA processing factors. Although elongation is stalled at the end of a short G-less cassette, time-course experiments show that CTD phosphorylations and associated factors continue to exchange as a function of time rather than location along the gene. Chemical inhibition shows that binding of Paf1C, capping enzyme, and Set2 to ECs requires TFIIH kinase (Kin28/Cdk7) activity. As this *in vitro* system recapitulates many known features of transcription elongation, it can be used to better understand factor dynamics as RNApII transitions from initiation to elongation, and how transcription is coordinated with nascent RNA processing and chromatin modifications.

## Results and Discussion

### Mass spectrometry analysis of RNApII ECs formed on immobilized templates

We sought to characterize RNApII ECs using the immobilized template assay and quantitative mass spectrometry approach (ITA/MS) previously used for studying PICs (Cho et al., 1997; Sikorski et al., 2012). The DNA template, fixed to magnetic beads via a biotin-streptavidin linkage, has five binding sites for the transcription activator Gal4-VP16 upstream of the *CYC1* core promoter (**Fig. 1a**). Because RNApII can elongate off the end of the linear template, a G-less cassette was used to stall ECs. PICs were assembled for 30 min in nuclear extract, and transcription then initiated by adding 3 NTPs (ATP, CTP, and UTP) and the chain terminator 3’-O-methyl-GTP (**Fig. 1b**). Transcription initiates within the G-less cassette just downstream of the promoter and extends until stalling where GTP should be incorporated (Hahn et al., 1985). After 15 minutes, the beads were washed extensively and candidate EC components eluted with the restriction enzyme SspI, which cuts just upstream of the transcription start site (TSS) to leave promoter-bound proteins on the beads (**Fig. 1a, b**).

**Figure 1.**
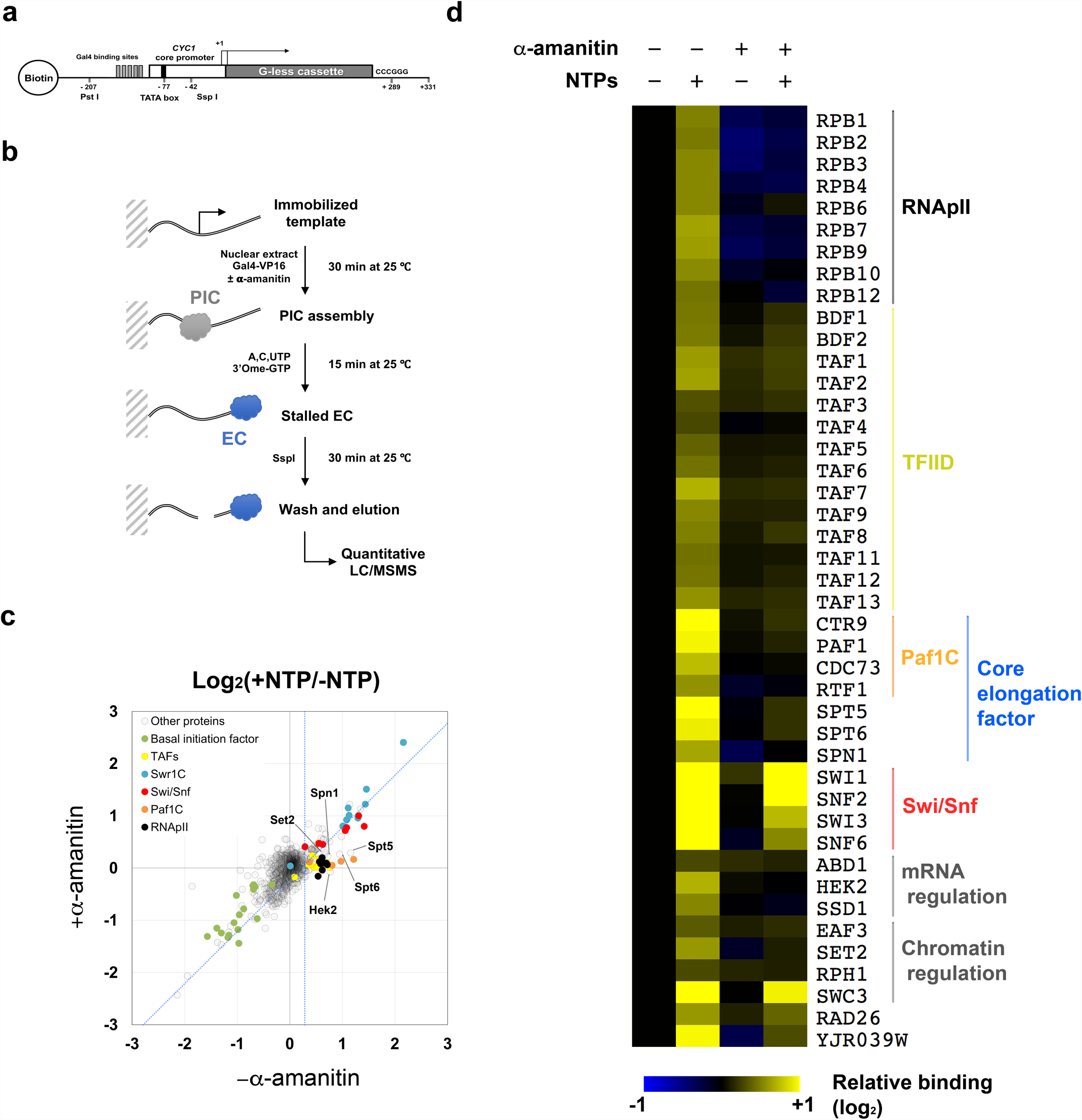
Characterization of RNApII elongation complexes generated *in vitro*. (**a**) Schematic diagram of DNA template used for RNApII EC isolation and analysis. (**b**) Experimental procedure for purification of RNApII ECs. PICs were pre-assembled on bead-immobilized DNA templates for 30 min in yeast nuclear extract with Gal4-VP16, either in the absence or presence of α-amanitin (1 µg/ml). Transcription was initiated by addition of NTPs (ATP, CTP, UTP and 3’-Ome-GTP). After 15 min of transcription, downstream bound proteins were released by Ssp I digestion and subjected to online 3D chromatography and LC-MS/MS with 4-plex iTRAQ reagents (Ficarro et al., 2011). **(c)** Log_2_ scaled +NTP/-NTP ratio of identified proteins were compared in the absence (x-axis) or presence (y-axis) of α-amanitin. Each circle represents a single protein, with the value calculated from the sum of all peptide signals assigned to the protein (see details in Materials and Methods). Color-coding is used to show members of specific complexes or functional subgroup as noted. Proteins specifically enriched by active transcription are defined as those to the right of the vertical blue-dotted line on the x-axis (95% probability for enrichment with NTPs) and below the diagonal blue-dotted line (95% probability for inhibition by α-amanitin). **(d)** Heatmap shows relative binding of transcription-dependent proteins. Signals were normalized by setting the-NTPs without α-amanitin channel to 1.0. See **Supplemental Fig. 1b and Supplemental Table 1** for additional data.

Samples were prepared with and without NTPs. To distinguish transcription from other NTP-dependent enzymatic reactions that may affect protein-DNA interactions (e.g. kinases, helicases, and other ATPases), the same experiment was carried out in parallel in the presence of the elongation inhibitor α-amanitin. Quantitative mass spectrometry using 4-plex iTRAQ labeling identified over one thousand proteins (**Fig. 1c, Supplemental Fig. 1a,** and **Supplemental Table 1**). Signals for the vast majority of proteins were the same under all conditions, thereby defining non-specific background binding. Against this null-hypothesis distribution we established a 95% confidence interval to identify three groups of proteins with distinct binding profiles (**Supplemental Fig. 1b**). Binding for the first group (22 total) was strongly increased by NTPs, regardless of whether transcription was inhibited (**Supplemental Fig. 1d**). These include the Swr1 Complex (Swr1C), RNApIII subunits, Snf5 and Swp82 of the Swi/Snf complex, the SUMO protein Smt3, Rtt107, multiple DNA repair proteins, and SSA related heat shock proteins. Binding of fifty-five proteins was reduced in the presence of NTPs, both with and without α-amanitin (**Supplemental Fig. 1c**). These include basal initiation factors for RNApII, negative cofactor 2 complex (NC2), Mot1, nucleotide excision repair factors, replication factors, mRNA binding proteins, and several sequence-specific transcription factors. The eluted downstream DNA has no consensus TATA, but some AT-rich sequences there probably support low levels of PIC formation. ATP hydrolysis may destabilize these non-productive PICs via the TFIIH translocase or by the Mot1 ATPase and NC2.

The final group of 43 proteins showed increased binding with NTPs, which was reduced by α-amanitin, indicating their presence on downstream DNA was transcription-dependent (**Fig. 1d** and **Supplemental Table 1**). This cluster contains all the RNApII subunits detected in this experiment (three small subunits were seen in later experiments described below), as well as known elongation factors Spt5/DSIF, Spt6, Spn1, and Paf1C. Therefore, we conclude that bona fide ECs are enriched in this *in vitro* system. The Rad26/CSB translocase was also in this group, consistent with its function pushing stalled ECs forward (Xu et al., 2017a). Unexpectedly, the TBP-associated factor (TAF) subunits of TFIID were found on downstream DNA, dependent on transcription. In a separate report we show these TAFs are not associated with ECs. Instead, TAFs interact post-transcriptionally with sequences just downstream of the TSS to promote re-initiation (Joo et al., 2017).

Although naked DNA templates were used in this experiment, several chromatin-related factors showed transcription-dependent enrichment. Histone chaperone Spt6 and the histone H3 K36 methyltransferase Set2 interact with phosphorylated Rpb1 (Li et al., 2007; Sdano et al., 2017; Yoh et al., 2007). The Eaf3 chromodomain protein, which binds methylated H3 K36 as part of the NuA4 and Rpd3S complexes (Li et al., 2007), and the H3 K36 demethylase Rph1were also isolated in this group. However, these were not reproducibly enriched in other EC experiments (see below). Finally, binding of four subunits of Swi/Snf (Swi1, Snf2, Swi3, and Snf6) is reduced by α-amanitin, although not to the same extent as RNApII or other EC components. While normally thought of as being targeted to promoters by activator, there is some evidence for Swi/Snf association with elongation (Schwabish and Struhl, 2007). Histones were detected in all four samples. Although these are often present as abundant contaminants in mass spectrometry-based experiments, we cannot rule out some chromatinization of the DNA templates in the extract (but see below).

Several RNA-binding proteins were identified as transcription associated factors. One is the mRNA cap methyltransferase Abd1. Unlike the capping enzyme guanyltransferase Ceg1 and triphosphatase Cet1, Abd1 crosslinks beyond promoters throughout transcribed regions (Komarnitsky et al., 2000; Schroeder et al., 2000). The hnRNP K protein Hek2/Khd1 binds many mRNAs through a CXXC motif (Hasegawa et al., 2008) that is also present in the G-less cassette. Hek2 co-immunoprecipitates with RNApII (Harlen et al., 2016). Also seen was the RNA binding protein Ssd1. Ssd1 binds CTD *in vitro* and shows genetic interactions with many transcription-related factors (Phatnani et al., 2004). Although these RNA binding proteins might interact with EC proteins, we think it more likely they bind the nascent transcript. The unusual base composition of the G-less transcript may explain the absence of better-known mRNA binding factors such as Npl3 and Hrp1. Future experiments with genes containing introns and polyadenylation sequences will be informative.

Compiled results from three biological repeats, each with two technical repeats, show the immobilized template approach reliably and reproducibly isolates promoter-driven RNApII ECs in nuclear extracts (**Supplemental Fig. 1e**). Quantitative mass spectrometry provides sensitive and accurate identification of EC-associated factors, even within a complex mixture. Thus, the ITA/MS approach is highly complementary to work with purified factors or *in vitro* analyses such as ChIP.

### RNApII CTD cycle progression is a function of time, not distance

The G-less cassette template used here stalls RNApII 289 bp downstream of the major proximal *CYC1* TSS (Hahn et a., 1985). Based on ChIP experiments (Komarnitsky et al., 2000; Mayer et al., 2010), we expected ECs isolated at this position to be in early stages, with high Ser5P and low Ser2P. Contrary to our expectations, identified proteins did not include early binding factors such as capping enzyme, but did include the CTD Ser2P-binding protein Set2. We realized that, while ChIP experiments measure EC association as a function of location on the gene, factor exchanges might reflect time elapsed post-initiation rather than the distance traveled.

To test if we had missed early-stage complexes, PICs were assembled on bead-immobilized templates as before, and ECs isolated over a time course of 0, 1, 2, 4, 8, and 16 minutes after NTP addition (**Fig. 2a**). CTD phosphorylation was monitored by immunoblotting with specific antibodies, probing either total bead-associated proteins (**Fig. 2b** upper panels, quantitated in **Fig. 2c**) or SspI-released downstream DNA (**Supplemental Fig. 2a**). As PICs form with non-phosphorylated polymerase (Laybourn and Dahmus, 1989), neither Ser5P nor Ser2P was detected before NTP addition (0 min). Remarkably, Ser5P increased sharply to its maximum level within the shortest time tested (1 min). After 2 minutes, it began to diminish until the last point assayed (16 min). Conversely, very low levels of Ser2P were detected one minute after the start of transcription and increased gradually, reaching maximum level at the final time point. Total Rpb1 binding declined slightly during this time. Therefore, yeast nuclear extracts can reproduce the progression of CTD phosphorylations seen *in vivo*.

**Figure 2.**
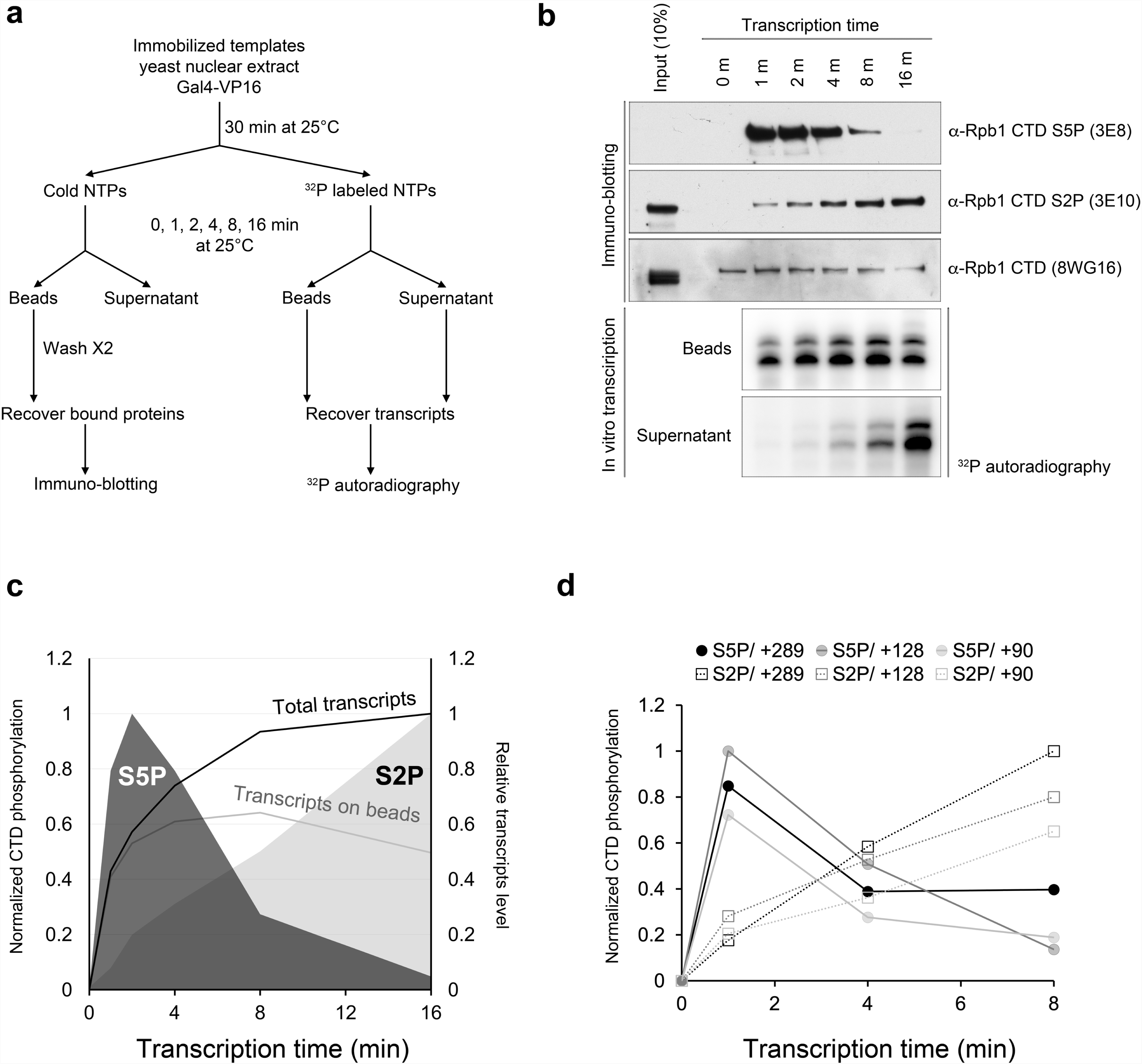
Time-dependent progression of CTD phosphorylation *in vitro*. (**a**) Experimental workflow of the time-course experiment for *in vitro* transcription and ITA. After PIC pre-assembly, samples were split in half. Transcription was started with NTPs (ATP, CTP, UTP and 3’-Ome-GTP). In one half reaction, ^32^P-labeled UTP was included for transcript detection. Proteins bound to the immobilized template were recovered magnetically after 0, 1, 2, 4, 8 and 16 min. (**b**) Samples with the non-labeled half reaction were immunoblotted with antibodies against “total” Rpb1 CTD (8WG16, which recognizes repeats not phosphorylated on Ser2), CTD Ser5P (3E8), or CTD Ser2P (3E10), with 10% of input extract used as a marker (upper panels). No signal was detected for Ser7P (4E12). With the ^32^P-labeled half reactions, nascent (Beads) and released (Supernatant) transcripts were recovered and detected by autoradiography (Lower panels). **(c)** Levels of the two CTD phosphorylations, normalized to total CTD, were quantitated and graphed as a function of time. Relative levels of total transcripts and nascent transcripts on beads were quantitated by phosphorimager densitometry and presented in the same graph. **(d)** CTD phosphorylations were measured after 0, 1, 4 and 8 min of *in vitro* transcription on different length G-less templates (90, 128, and 289 nt) as in parts **b** and **c**. See **Supplementary Fig. 2b** for additional information.

To demonstrate that these changes occur in stalled ECs, gel electrophoresis was used to analyze radioactive transcripts retained on beads or released into the supernatant (**Fig. 2b** lower panels, quantitated in **Fig. 2c**). The bulk of full-length transcription was observed within the first 4 minutes, with a slow increase thereafter that may be due to later initiation events. Binding of the EC at the stall site was stable for up to 8 min. Some transcript release was seen at 4 and 8 minutes, and was quite appreciable by 16 minutes, when Ser2P was highest. The kinetics of CTD phosphorylation changes were similar when ECs were stalled at +90, +128, or +289 (**Fig 2d**, **Supplemental Fig 2b**), but not when PICs were treated with ATP alone (**Supplemental Fig 2c**).

These results lead to several conclusions. First, the CTD phosphorylation cycle, previously inferred from ChIP experiments, can be reproduced *in vitro*. This system will facilitate future studies of the relevant kinases and phosphatases. Second, the progression from Ser5P to Ser2P continues despite the lack of forward movement in stalled ECs. This finding complements ChIP experiments showing that RNApII mutants with accelerated or slowed polymerization cause CTD phosphorylation peaks to shift slightly downstream or upstream *in vivo*, respectively (Fong et al., 2017; Soares et al., 2017). These results argue that the CTD cycle is generated as a function of time rather than distance from the start of transcription.

### Temporally regulated exchange of EC components

To analyze changes in EC components, ITA/MS was performed over a similar time course (**Fig. 3a**). Over a thousand proteins were identified on the downstream template after NTP addition, with most showing no significant changes (**Supplemental Fig. 3a**). As in **Fig. 1**, binding of the basal transcription factors, NC2, and Mot1 decreased (**Supplemental Fig. 3c**). In contrast, addition of NTPs increased binding of roughly 10% (105) of detected proteins (**Supplemental Fig. 3b**). Time-dependent changes in NTP-enriched proteins were quantified by calculating the slope of a line through their post-NTP time point values (**Supplemental Table 1**, shown as green/pink scale in **Figs. 3 and 4**).

**Figure 3.**
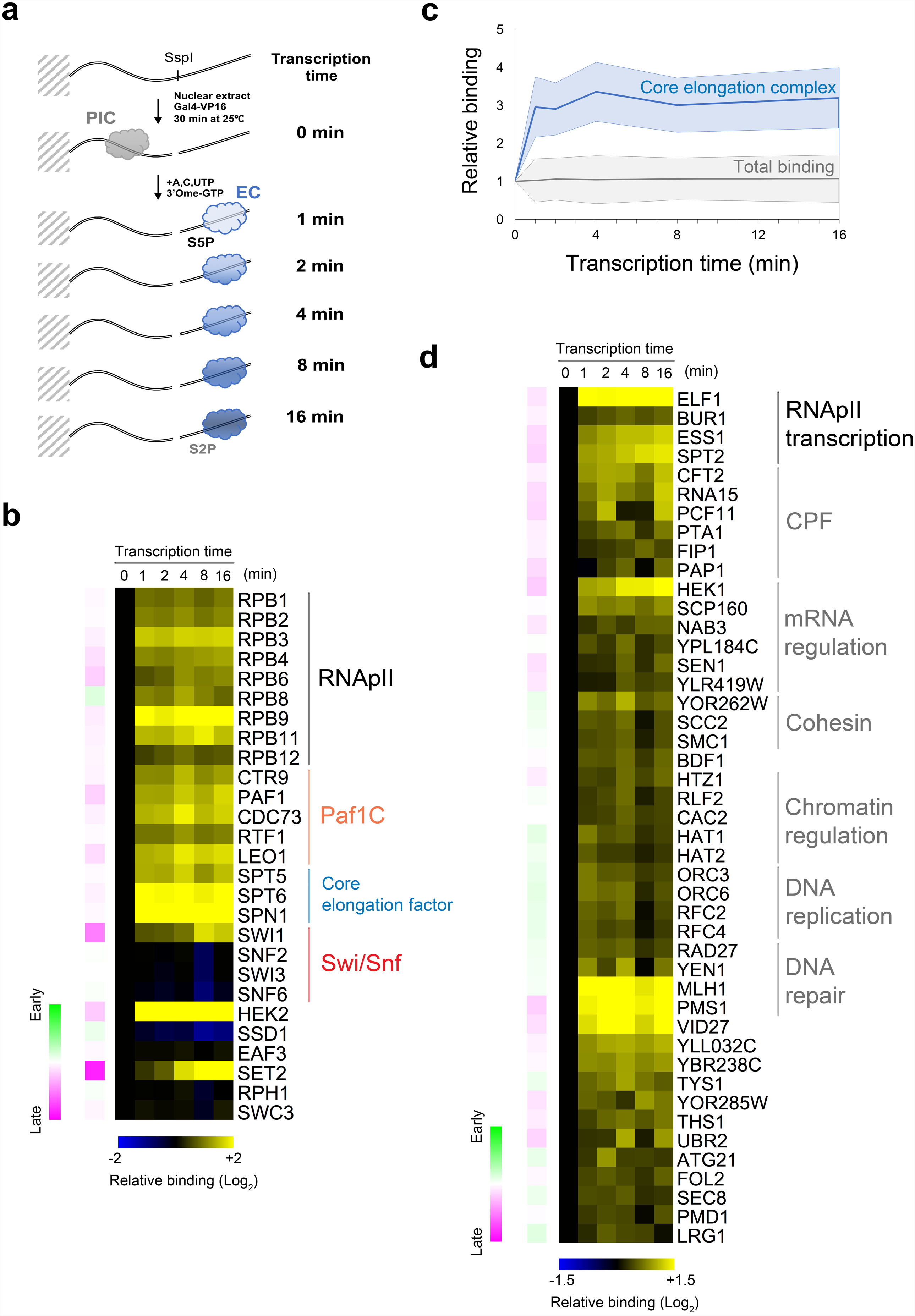
Time course analysis showing EC factor exchange. (**a**) Schematic diagram of the time-course experiment. After preassembling PICs, transcription was started by NTPs (A, C, UTP and 3’-Ome-GTP). Downstream bound proteins were eluted by SspI digestion at 0, 1, 2, 4, 8, or 16 min after transcription start. Eluted proteins were analyzed by 6-plex TMT-labeled quantitative 3D LC-MS/MS. (**b**) Heatmap shows protein levels relative to the 0 time point (blue/yellow scale), and slope of change for the 1 - 16 minute time points (pink/green scale, with pink signifying an increase and green a decrease over time). **(c)** Normalized and averaged signals for RNApII subunits and core elongation factors are graphed (blue line) with standard deviation (blue shading), compared to total identified proteins (gray line and shading). **(d)** Heatmap shows relative levels and slope of change for additional proteins significantly and consistently enriched downstream by NTPs. These proteins are defined as having greater than 95% probability for enrichment by NTPs (averaged for 1, 2, 4, 8, and 16 min time points) but below 95% probability for significant slope change compared to all other proteins. See **Supplemental Figs. 3b, c and Supplemental Table 1** for additional data.

**Figure 4.**
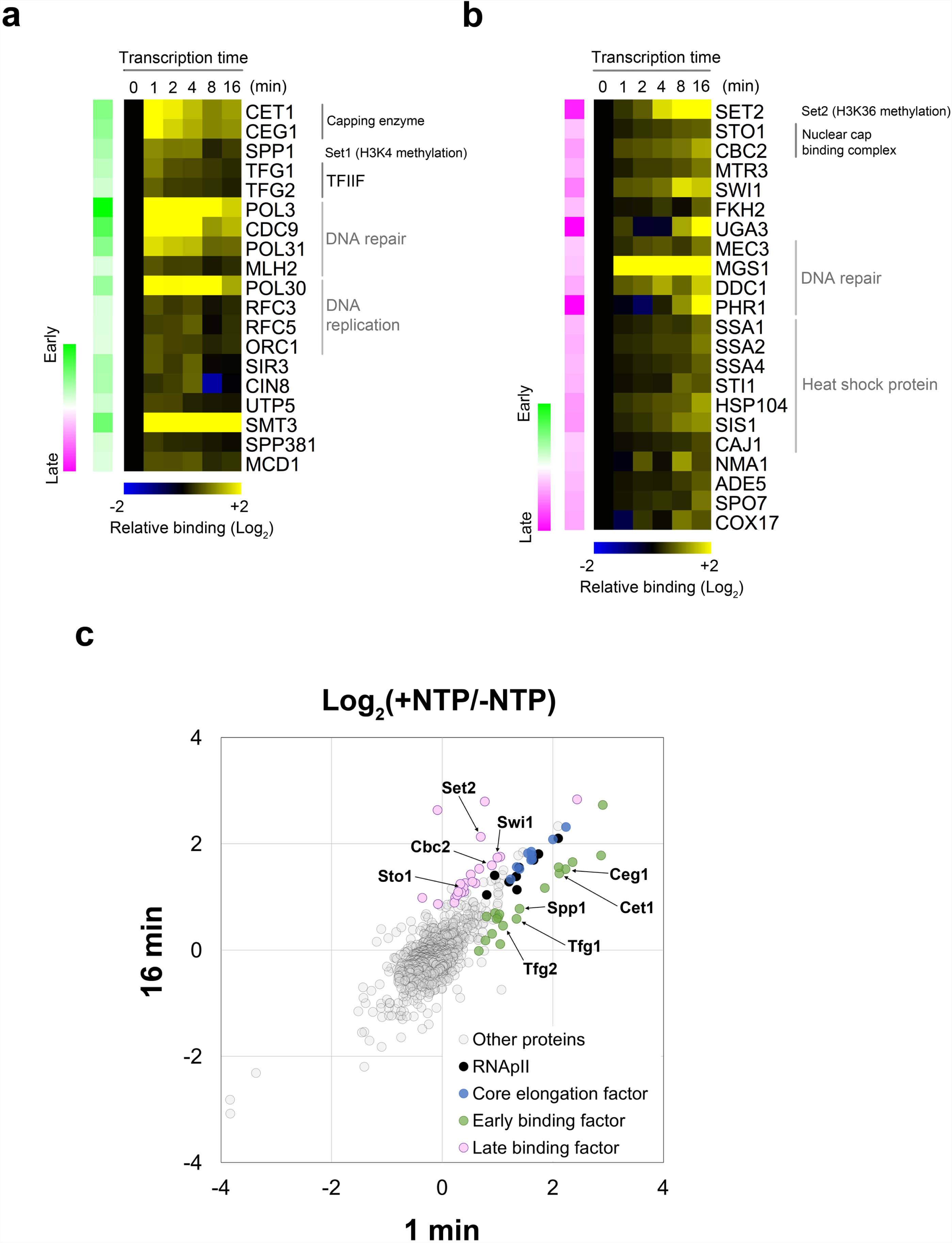
EC-associated factors that change over the time course. Heatmaps show relative protein levels and slope of change for (**a**) early-enriched (strong positive slope, green) and (**b**) late-enriched (strong negative slope, pink) factors from the quantitative mass spectrometry experiment shown in **Fig. 3**. Differential binding factors are defined as those having 95% probability for being enriched by NTPs and 95% probability for the slope being outside the distribution of slopes for all proteins. **(c)** Scatter plot showing log_2_ scaled ratio for total proteins at the early (1 min on x-axis) and the late (16 min on y-axis) time points relative to 0 time point. Each circle represents a single protein. The early (green) and late (pink) binding factors shown in panel (**a**) and (**b**) are color-coded along with RNApII (black) and core elongation factors (blue). See also **Supplemental Fig. 3b** and **Supplemental Table 1**.

RNApII subunits appeared downstream within one minute and remained steady during the entire reaction (**Figs. 3b, c**), consistent with levels of nascent transcripts on the beads (**Figs. 2b, c**). Known elongation factors Paf1C, DSIF(Spt4-Spt5), Spt6-Spn1, and Elf1 were also rapidly and strongly recruited by NTPs, and the ratio of these factors to RNApII remained constant throughout the reaction (**Figs. 3b, c**). TFIIS (Dst1) also increased after NTP addition, but not above the level of statistical significance. Structural studies show these factors bind directly to RNApII (Ehara et al., 2017; Vos et al., 2018; Xu et al., 2017b). They also crosslink throughout transcribed regions in ChIP studies, and mutations cause phenotypes associated with elongation defects. Based on these criteria, we designate this group of EC-associated proteins as “core elongation factors”.

Other factors that showed increased binding at all time points include the CTD prolyl isomerase Ess1, the Cdk9 homolog Bur1/Sgv1, and Spt2/Sin1, all thought to function during elongation (**Fig. 3d**). We did not detect the CTD Ser2 kinase Ctk1/Cdk12, nor the CTD phosphatases Ssu72, Rtr1, or Fcp1. As these are presumably required for progression of CTD phosphorylations, these proteins may be below the level of detection, or like many kinases and phosphatases, may bind their substrates only transiently. Addition of NTPs also increased levels of the SUMO protein Smt3 and many DNA repair factors. However, their binding was not sensitive to α-amanitin and therefore independent of transcription (**Fig. 1**). Interestingly, although FACT (Spt16-Pob3), Chd1, SAGA, the ISWI complexes, Def1, and Elongator have been implicated in elongation, and many peptides from these factors were detected, none of them showed EC enrichment on naked DNA templates.

Multiple RNA binding factors were again enriched upon NTP addition, despite non-specific competitor RNA present in the reaction. Most strongly recruited were the KH-domain proteins Hek2 and Pbp2/Hek1, the yeast homologs of hnRNP K protein. A third KH-domain protein, Scp160, was also enriched, as was Mrn1/YPL184C, a protein implicated in mRNA maturation. As in **Fig 1**, known mRNA binding proteins such as Hrp1, Npl3, Yra1, Mex67, or THO/TREX were not enriched, suggesting the ECs generated *in vitro* on this template lack the appropriate RNA sequences or some other factor needed for their stable binding.

Other mRNA processing factors enriched by NTPs include multiple cleavage and polyadenylation factors (**Fig. 3d**). Lacking a polyadenylation site, there is no apparent cleavage of the G-less RNA transcript (**Fig. 2b**), and no enrichment of the “torpedo” termination factors Rtt103, Rat1, or Rai1 was seen. The Nab3 and Sen1 proteins, which cooperate with Nrd1 to terminate transcription on snoRNAs and other small non-coding transcripts (Steinmetz et al., 2001), scored as EC-associated in this experiment (**Fig. 3d**). Splicing factors detected were not enriched by NTPs, with the possible exception of Spp381, a U4/5/6 subunit that showed weak early binding. In the future it will be informative to analyze ECs on templates containing introns, polyadenylation sites, or Nrd1/Nab3 terminators.

Some proteins showed strong differential binding over time, as calculated from the time course slope (**Fig. 4c, Supplemental Table 1**). For example, mRNA capping enzyme (a heterodimer of the triphosphatase Cet1 and guanylyltransferase Ceg1) showed strongest binding immediately after addition of NTPs (**Fig. 4a**, 1 min). Previous immobilized template experiments showed that ATP alone triggers CTD phosphorylation and capping enzyme association with PICs (Cho et al., 1997). Paralleling Ser5P (**Fig. 2b**), capping enzyme association gradually decreased.

Several other proteins related to gene expression showed strong binding to early ECs (**Fig. 4a**). In marked contrast to other basal initiation factors, TFIIF (a heterodimer of Tfg1 and Tfg2) was enriched at early time points. TFIIF affects promoter clearance, can bind RNApII ECs, and stimulates elongation *in vitro* (Thomas and Chiang, 2006). However, ChIP experiments do not detect TFIIF crosslinking in transcribed regions (Rhee and Pugh, 2012). Our results suggest TFIIF may remain bound to RNApII during very early elongation. RNA polymerases I and III have TFIIF-like integral subunits that presumably remain associated throughout transcription (Vannini and Cramer, 2012), so dissociation of TFIIF soon after promoter escape may be a RNApII-specific feature.

Also showing early binding is the Spp1 subunit of the Set1/COMPASS H3K4 histone methyltransferase, which binds Ser5P-associated CTD. Other COMPASS subunits were not detected in this experiment, so the significance of this finding remains unclear. The Sub1/PC4 protein, which associates with PICs and perhaps the transcription bubble (Sikorski et al., 2011), was observed at early time points just below the statistical cutoff for 95% confidence.

Factors preferentially binding late ECs included Cap Binding Complex (CBC, a heterodimer of Sto1 and Cbc2) and the histone methyltransferase Set2 (**Fig. 4b**). Set2 directly binds Ser2P CTD, explaining the increase over time. CBC is not known to directly bind CTD, so interaction is mostly likely with the newly formed cap structure of the nascent mRNA (derived from the 3’-Ome-GTP in the reaction).

Multiple experiments suggest CBC functions in ECs: CBC ChIPs to transcribed regions, and strong genetic interactions of CBC genes with Set2 and other elongation factors have been reported (Hossain et al., 2013). Late binding was also seen for Fkh2, another protein implicated in elongation (Morillon et al., 2003). The exosome subunit Mtr3 also showed a significant increase over time, although other subunits of the complex were not enriched by NTPs. Several heat shock and DNA repair proteins also increased, but their enrichment was also seen in the absence of transcription (**Fig. 1**).

### Kin28/Cdk7 kinase activity is required for both Ser5 and Ser2 phosphorylation

The time-course experiment showed clear correlations between CTD phosphorylations and differential EC binding of several factors. To directly test the role of CTD phosphorylation, we used the Kin28is double mutant (L83G, V21C), which can be irreversibly inactivated by the chemical inhibitor CMK (Rodriguez-Molina et al., 2016). As previously reported, CMK strongly inhibited growth of Kin28is, but not wild-type, cells (**Supplemental Fig. 4a**). Upon treatment of Kin28is nuclear extract with CMK, *in vitro* phosphorylation of Ser5 was effectively blocked (**Fig. 5a, Supplemental Fig. 4b**).

**Figure 5.**
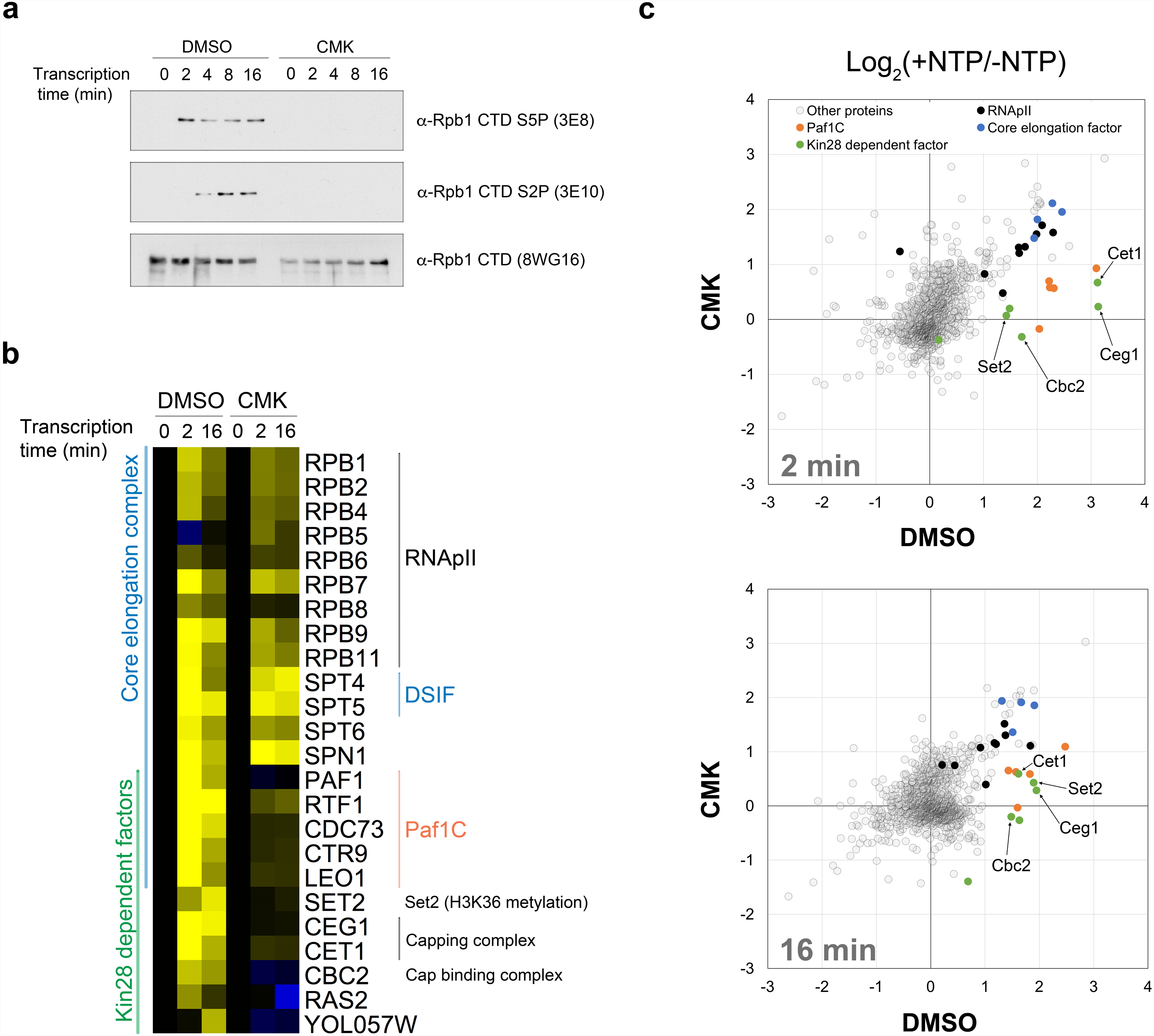
Kin28/Cdk7 activity is required for CTD phosphorylation and association of a subset of EC factors. (**a**) PICs were assembled in Kin28is nuclear extract in the presence of CMK inhibitor (250 nM) or DMSO solvent. Total bound proteins were recovered 0, 2, 4, 8, and 16 min after transcription initiation with NTPs (A, C, UTP and 3’-Ome-GTP). CTD phosphorylation was monitored with specific antibodies against CTD Ser5P (3E8), CTD Ser2P (3E10), and “total” CTD (8WG16). (**b**) Reactions were performed as in panel **a**, except proteins bound to template were eluted by PstI digestion at 0, 2, or 16 min after transcription start. Proteins were subjected to 6-plex TMT-labeled quantitative 3D LC-MS/MS analysis. The heatmap shows relative levels for RNApII subunits, core elongation factors, and Kin28-dependent factors. Kin28-dependent factors were defined as those with at least 95% confidence level for enrichment by NTPs (log_2_ of the 2 or 16 min +DMSO signal divided by the zero time point signal) and also a 95% confidence level for significant reduction upon CMK treatment (log_2_(2 or 16 min/0 min) with DMSO – log_2_(2 or 16 min/0 min) with CMK inhibition). Note that Ras2 and YOL057W did not score as transcription-enriched in other experiments and are likely false positives. **(c)** Scatter plots showing log_2_ scaled +NTP/-NTP ratios of 1,240 identified proteins in the presence of DMSO (x-axis) versus CMK (250 nM; y-axis), either during early elongation (2 min, upper panel) or late elongation (16 min, lower panel). Colors designate RNApII subunits (black), core elongation factors (blue), Paf1C subunits (orange), and other Kin28-dependent factors (green). See also **Supplemental Figure 4 and Supplemental Table 1**.

Surprisingly, Ser2P was also completely blocked by Kin28 inhibition. This was unexpected, as *in vivo* inhibition of Kin28/Cdk7 in budding or fission yeast cells is reported to have little effect on Ser2P (Bataille et al., 2012; Booth et al., 2018; Mbogning et al., 2015; Rodríguez-Molina et al., 2016; Tietjen et al., 2010). By comparison, both Ser5P and Ser2P were only partly reduced upon Cdk7 inhibition in mammalian cells carrying a mutant kinase sensitized to a non-covalent inhibitor (Ebmeier et al., 2017; Larochelle et al., 2012), perhaps due to incomplete kinase inactivation. In contrast, the covalent Cdk7 inhibitor THZ1 strongly blocked both Ser5 and Ser2 phosphorylation *in vivo* (Kwiatkowski et al., 2014) and *in vitro* (Nilson et al., 2015), but this chemical also inhibits the Ser2 kinase Cdk12.

Loss of Ser2P upon Kin28is inactivation is not due to absence of transcription (**Fig. 6a**). It is unlikely that Kin28 directly phosphorylates Ser2, given its preference for Ser5 (Eick and Geyer, 2013; Komarnitsky et al, 2000). Presumably, Kin28 activity on the CTD or some other substrate promotes Ser2 phosphorylation by another kinase. *In vitro*, yeast Ctk1 (Jones et al., 2004), metazoan Cdk12 (Bartkowiak and Greenleaf, 2014), and mammalian Cdk9 (Eick and Geyer, 2013) more efficiently phosphorylate Ser2 when CTD is “primed” by prior phosphorylation at Ser5 or Ser7. In mammalian cells, Cdk7 directly phosphorylates and activates Cdk9 (Larochelle et al., 2012). Finally, Ser5P may recruit one or more Ser2 kinases (see below). Although the mechanism remains unclear, the requirement of Kin28 activity for subsequent Ser2 phosphorylation helps explain the sequential nature of CTD modifications.

**Figure 6.**
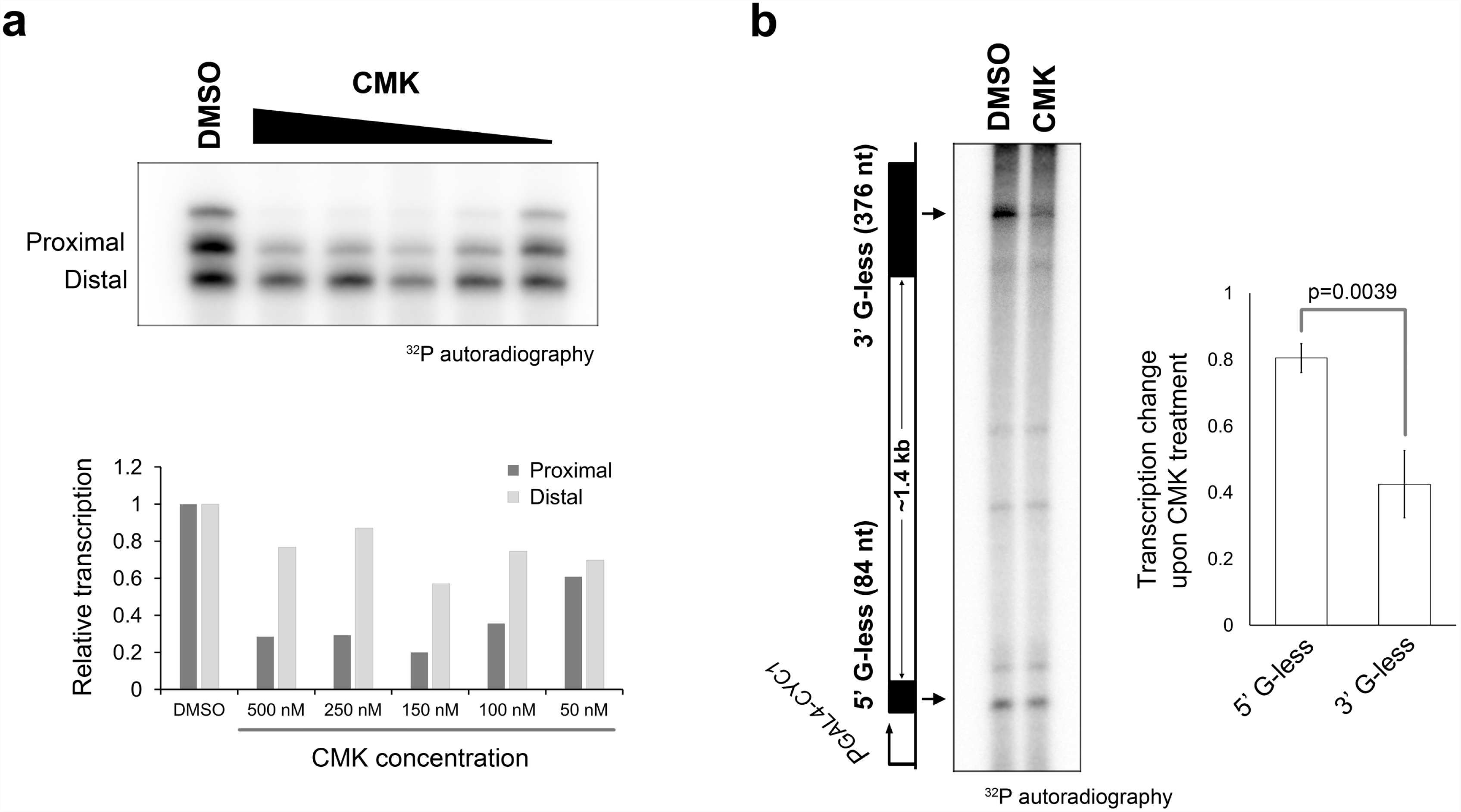
Kin28 kinase inactivation affects transcription start site selection and elongation processivity. (**a**) Kin28is nuclear extract was preincubated with the indicated concentrations of CMK for 10 min, followed by *in vitro* transcription on template pUC18-G5CYC1 G-(SB649). Labeled transcripts were separated on a 6% urea-acrylamide and detected using autoradiography (upper panel). The lower panel shows normalized quantitation of transcripts from proximal and distal start sites. Note that the uppermost band results from read-through transcription through the full G-less cassette. (**b**) *In vitro* transcription was performed with KIN28is nuclear extract, pre-incubated with 500 nM CMK or DMSO solvent. The template was pGCYC1-402 (F647, Rondon et al., 2003), which produces a 1.4 kb transcript containing two different length G-less cassettes, one near the 5’ end and one near the 3’ end. After RNAse T1 digestion, resistant fragments were separated on a 6% urea-acrylamide gel (left panel). Relative change upon CMK treatment for each fragment is shown in right panel. Error bars designate standard deviation in three independent reactions, calculated using the Student t-test. See also **Supplemental Figure 5.**

### Kin28/Cdk7 kinase activity is required for recruiting a subset of elongation complex components

To identify which EC components are dependent on CTD phosphorylation, Kin28is nuclear extract was treated with CMK or solvent (DMSO) for 30 minutes during PIC formation on immobilized templates. ECs were recovered by PstI digestion at 0, 2, and 16 minutes after addition of NTPs, followed by multiplexed quantitative mass spectrometry (**Figs. 5b, c, Supplemental Table 1**). NTP-dependent stimulation of RNApII and core elongation factors Spt4-Spt5, Spt6, and Spn1 remained high. Only a limited set of EC factors were strongly reduced upon Kin28is inhibition (**Figs. 5b, c**). The two mRNA capping enzyme subunits and Set2 topped the list, as expected from their interactions with Ser5P and Ser2P, respectively. CBC was also reduced, presumably due to the lack of capping. Interestingly, Spt6 was unaffected, consistent with recent data suggesting it binds to the Rpb1 linker rather than the CTD (Sdano et al., 2017; Yoh et al., 2007). Finally, all five subunits of Paf1C were strongly reduced by Kin28 inhibition.

The drop in Paf1C was unexpected because Paf1C ChIP is independent of Ser2P phosphorylation by Ctk1, yet its crosslinking pattern does not match Ser5P (Ahn et al., 2004). Furthermore, recent structures show Paf1C interacting with the body of RNApII, with no apparent CTD contacts (Ehara et al., 2017; Vos et al., 2018; Xu et al., 2017b). Paf1C can bind phosphorylated CTD *in vitro* (Phatnani et al., 2004; Qiu et al., 2012), but Paf1C recruitment to ECs is thought to primarily involve binding to an Spt5 domain that is phosphorylated by Bur1/Cdk9 early in elongation (Liu et al., 2009; Zhou et al., 2009). Qiu et al. (2012; 2009) showed this Spt5 phosphorylation is blocked by Kin28 inhibition, and suggested this effect was due to Bur1 directly binding CTD Ser5P. However, Bur1-Bur2 association with ECs was not blocked by CMK in our experiment (**Supplemental Table 1**). Therefore, how Kin28 activity promotes Paf1C binding remains an open question.

Loss of Paf1C from the EC is predicted to affect elongation. Indeed, Kin28 inhibition dramatically reduced the ratio of distal to proximal RNA on a long *in vitro* transcription template (**Fig. 6b**). Paf1C loss may also explain the elongation defect seen upon *in vivo* inhibition of Kin28is (Rodríguez-Molina et al., 2016; Booth et al., 2018). Interestingly, the small drop in transcription upon CMK treatment, seen with Kin28is but not wild type nuclear extract, primarily reduced the proximal *CYC1* TSS (**Fig. 6a, Supplemental Fig. 5a, b**). Similar TSS effects are seen with certain TFIIB mutants and RNApII mutants that slow elongation rate (Kaplan et al., 2012). We note that Murikami et al. (Murakami et al., 2015) also reported a role for Kin28 in determining TATA box-TSS spacing, but that function is clearly distinct, as it shifts TSSs downstream and is kinase independent. Future experiments will explore the roles of Kin28 kinase activity in TSS selection.

## Conclusions

Using quantitative proteomics to analyze RNApII complexes on immobilized template DNA, we show that yeast nuclear extract reproduces many key aspects of elongation, including CTD phosphorylation changes and dynamic exchange of associated factors. We identified both “core elongation factors” that track with RNApII levels at all time points (**Figs. 1 and 3**) and other EC proteins that change over time (**Fig. 4**). Remarkably, ECs continue progression through the CTD cycle even when elongation is stalled (**Fig. 2**), demonstrating that these transitions as a function of time, rather than distance traveled, after initiation.

The ITA/MS approach will be very useful for analyzing the dynamics of RNApII elongation. The ability to isolate and characterize *in vitro* ECs at various post-initiation times provides a powerful system for studying the interplay between CTD modifications, elongation factors, and the chromatin template.

## Acknowledgements

This work was supported by DFCI/Blaise Proteomics Center support to J.M. and NIH grants GM46498 and GM56663 to S.B.

## Author Contributions

All experiments were performed by Y.J., with S.B.F. also contributing to mass spectrometry data collection and analysis. Y.C. constructed the Kin28is strain and provided other technical support. Y.J., S.B.F., J.A.M., and S.B. designed the experiments. Y.J. and S.B. wrote the manuscript.

## Competing Interests

The authors declare no competing interests.

## Materials and Methods

### Yeast Strains, plasmids and oligonucleotides

Yeast strains, oligonucleotides and plasmids used this study are listed in **Supplemental tables 2, 3, 4**.

### Yeast nuclear extract preparation and immobilized template binding assay (ITA)

Nuclear extracts were prepared as described previously (Sikorski et al., 2012) from wild type (BY4741 or YF702) or Kin28is (YSB3566) yeast strains. Immobilized template binding assays for ECs were performed as described for PICs (Sikorski et al., 2012), with modifications described below. Biotinylated DNA templates (**Fig. 1a**) were amplified by PCR from pUC18-G5CYC1 G-(SB649) with primers O#3151 and O#1477. Naked templates (200 ng per 1x reaction) were immobilized onto 50 μg /5 μl Dynabeads Streptavidin T1 slurry (Invitrogen) in 1X TE (10 mM Tris-HCl pH 8, 1 mM EDTA) supplemented with 1 M NaCl for one hour at 25°C with rotation. Chromatinized templates were similarly linked to Dynabeads, except in Binding Buffer (15 mM HEPES pH 7.5, 150 mM NaCl, 4% PEG 8000, 10 mM EDTA, 0.02% NP40, 5% Glycerol, 1.25 mM β-glycerophosphate, 0.1 mM PMSF, 0.5 mM DTT) for one hour at 25°C with rotation. To reduce background binding, template-coupled beads were blocked for 30 min at 25°C with rotation in 50 μl Transcription Buffer (20 mM HEPES-KOH pH 7.6, 100 mM potassium acetate, 1 mM EDTA, 5 mM magnesium acetate) complemented with 60 mg/ml casein, 5 mg/ml polyvinylpyrrolodone, and 2.5 mM DTT. After washing three times in 400 μl Transcription Buffer, the immobilized templates were resuspended in 10 μl Transcription Buffer. Recombinant transcription activator Gal4-VP16 (400 ng) was pre-incubated with immobilized templates for 5 min at 25°C prior to the addition of yeast nuclear extract. Extract (∼1 mg) and remaining components (20 units Rnasin, 2 units creatine phosphokinase, 10 mM phospho-creatine, 0.03% NP-40, 16.67 μg/ml tRNA, 160 µM S-adenosylmethionine, 20 µM acetyl-CoA, and 16.67 μg/ml HaeIII-digested *Escherichia coli* genomic DNA) were added to a total reaction volume of 60 μl. After 30 min incubation at 25°C to form PICs, transcription was initiated by adding 2.4 µl NTPs mix (final concentration of 400 µM each ATP, CTP, and UTP, and 40 µM 3’-Ome-GTP) to the mixture, followed by an incubation at 25°C with occasional gentle mixing every 5 min to keep beads suspended. After the time indicated, immobilized templates were recovered with a magnetic stand and the reaction was stopped by quickly washing twice in 200 μl of Wash Buffer (Transcription Buffer complemented with 0.05% NP40 and 2.5 mM DTT). For analysis of total bound proteins, samples were recovered by boiling with 40 µl Transcription Buffer and 40 µl 2X SDS loading dye (100 mM Tris-HCl pH 6.8, 4% SDS, 0.2% Bromophenol Blue, 20% Glycerol, 200 mM DTT). To analyze proteins bound to specific regions on the DNA template, complexes were eluted for 30 min at 25° with rotation in 40 μl of the Transcription Buffer with 60 units of either SspI-HF or PstI-HF (New England Biolabs) as indicated (see **Fig. 1**).

### Sample preparation for Quantitative mass spectrometry

Quantitative mass spectrometry was performed as described previously with minor modifications (Sikorski et al., 2012). Reactions were the same as described above for ITA, but scaled up 5-fold (300 µl reaction volume and 200 µl of final elute). EC formation was validated by immunoblotting 40 µl of the final restriction enzyme eluate with specific antibodies against Rpb1-CTD (8WG16), Rpb1-CTD Ser5P (3E8) and Rpb1-CTD Ser2P (3E10). The remaining eluate (160 µl) was processed for mass spectrometry. Ammonium bicarbonate was added to a final concentration of 50 mM to each sample. Protein cysteines were reduced with 10 mM DTT for 1hr at 56°C, and then alkylated with iodoacetamide (55 mM for 1 hour at 25°C in the dark). Trypsin was added at 1:20 (enzyme:substrate) ratio. After incubation at 37°C overnight, samples were acidified with trifluoroacetic acid (TFA) to pH ∼2 (measured by pH strip) and microfuged at 13,000 rpm to remove insoluble particles. Tryptic peptides were desalted and purified on a C18 solid phase extraction cartridge, eluted with 80% acetonitrile/0.1% TFA, and dried. For 4-plex iTRAQ reagent, each dried peptide pellet was resuspended in 30 μL 0.5 M triethylammonium bicarbonate and then 70 μL 100% ethanol was added. Each sample was mixed with a specific iTRAQ label and incubated for 1 hour at room temp. For 6-plex TMT labeling, each dried pellet was resuspended in 100 μL 0.5 M triethylammonium bicarbonate. Each sample was then mixed with a specific TMT reagent reconstituted in 40 μL acetonitrile, followed by incubation for 1 hour at room temp. After labeling, samples were combined and desalted with a C18 solid phase extraction cartridge and dried. Final peptide pellet was reconstituted in 80 µl 10 mM ammonium formate. Labeling efficiency above 95% was confirmed by 1D reverse phase LC-MS screening before online 3D chromatography and LC-MS/MS. For each technical repeat, half of each sample was injected. To estimate reproducibility across biological replicates, +/-NTP samples (15 minutes, naked DNA template) from three independent experiments (representing three biological replicates, each with two technical replicates) were analyzed. Over 700 proteins were identified in all six replicates (**Supplemental Table 5**). For each protein, the P-value for the six replicates (three biological times two technical repeats) falling inside the normal distribution for the entire dataset was plotted against the +NTP enrichment score, showing very strong statistical significance for enrichment of elongation factors and the large majority of RNApII, Paf1C, and TFIID subunits (**Supplemental Fig. 1e**).

### Online 3D Chromatography and LC-MS/MS Data Acquisition

The 3-dimensional chromatography system used for peptide separation was essentially as described (Ficarro et al., 2011; Zhou et al., 2011). Our RP-SAX-RP (reverse phase-strong anion exchange-reverse phase) platform is based on a Waters NanoACQUITY UHPLC system, equipped with a third, 6-port, 2-position valve (VICIValco, Houston, TX). True nanoflow rates in the third dimension were achieved through use of a passive split located prior to the third dimension PC. The valve then switches to engage the passive split such that a majority of LC effluent is diverted to waste prior to the third dimension PC. 13 RP-SAX-RP fractions were analyzed. Either an LTQ Velos (**Figs. 1, 7**) or Q-Exactive HF (Thermo Scientific) machine was used to acquire MS/MS spectra in data-dependent mode as described (Zhang et al., 2009) using the following parameters: MS spectra were acquired with a target value of 2 × 10^6^, maximum ion injection time of 500 ms, and a mass resolution of 60,000 (FWHM @ m/z = 400). A minimum threshold of 1 × 10^5^ was used for MS/MS, with a charge state range of 2+ to 4+. Up to eight precursors could be selected for each cycle, with a 20 s exclusion time for each precursor selected for MS/MS. One linear trap CAD and one HCD MS/MS spectrum were acquired in series for each precursor. An optimized collision energy was used for HCD scans.

MS/MS data files were directly accessed and converted to.mgf files with ytraq4plex.mz to extract CAD and HCD scans into separate files for subsequent MASCOT searches (Zhang et al., 2009). In addition, iTRAQ and TMT reporter ion peak intensity values were extracted from HCD scans, and then inserted in the header for corresponding CAD and HCD scans of the same precursor; this provided for convenient retrieval of both peptide sequence identification and reporter ion quantification data from the MASCOT search results. The two files were searched using Mascot version 2.3.1 against a yeast database from the Saccharomyces Genome Database (downloaded April 2010), with an appended cRAP (common repository of adventitious proteins) database. The mass tolerance values were set to 10 ppm for each precursor, 0.6 Da for LTQ-CAD scans, and 25 mmu for HCD scans. After searching, an Excel spreadsheet containing the Mascot search results was generated using Multiplierz software (Askenazi et al., 2009; Parikh et al., 2009). Scans with FDR <1% were extracted for further analysis, and HCD and CAD results were compressed into a single worksheet. For relative quantitation, iTRAQ or TMT reporter ion peak intensity values were extracted from HCD scans, corrected for known isotopic impurities, and included in the Multiplierz spreadsheet. The iTRAQ or TMT reporter signal intensity values of all peptide scans for a given protein were summed prior to calculation of ratios. Only peptides that were unique to the matched protein and detected in both replicates were used for quantitation. Detailed data and pre-normalized iTRAQ/TMT values for all individual peptides in each replicate are in **Supplemental Table 6**. The total number of peptide sequence matches was used to normalize each channel, with restriction enzyme or bovine serum albumin peptides from the restriction buffer acting as a “spike-in” control. Normalized data was statistically analyzed using the Mixed Model analysis within the R software package. Using a null hypothesis of no differences between conditions, ratios for all proteins are fit to a normal distribution. The Mixed Model calculates confidence level for individual protein ratios being outside the null hypothesis distribution.

### In vitro transcription

For *in vitro* transcription on immobilized templates shown in **Fig. 2b**, Gal4-VP16 (400 ng) and bead-linked DNA (200 ng) were mixed with ∼1 mg of yeast nuclear extract using the same conditions as for the immobilized template binding assay. For transcript detection, ^32^P-labeled NTPs (500 μM ATP, 500 μM CTP, 20 μM UTP, 50 μM of 3’-Ome-GTP and 3 μCi ^32^P-UTP) were added. After the time indicated, beads were separated from the supernatant on a magnetic stand. RNA was recovered from both fractions and detected on 8M urea - 6% polyacrylamide gel. *In vitro* transcription on plasmids was performed as previously described (Sikorski et al., 2012). After 15 min pre-incubation of CMK with 100 ng template plasmid, 200 ng Gal4-VP16, and ∼500 μg yeast nuclear extract, transcription is initiated by adding ^32^P-labeled NTPs (500 μM ATP, 500 μM CTP, 20 μM UTP, 50 μM of 3’-Ome-GTP and 3 μCi ^32^P-UTP) for 30 min at 25°C. For the *in vitro* elongation assay shown in **Fig. 6b**, pGCYC1-402 (F647) plasmid was used (Rondon et al., 2003). For the CMK titration assay in **Fig. 6a,** pUC18-G5CYC1 G-(SB649) plasmid was used.

