## Supplemental Figures 1-5 for "In vitro analysis of RNA polymerase II elongation complex dynamics"

Supplemental Figures 1-6

Supplemental Tables 1-7

**Supplemental Figure 1**, related to Figure 1**.** ITA/MS analysis of RNApII elongation complexes.

(**a**) Heatmap shows binding profiles for all identified proteins (1,511 total) from quantitative mass spectrometry. Relative reporter intensity of each condition was calculated by setting the untreated sample (-NTPs/-$\alpha$-amanitin) to 1.0. Each line represents one protein. Based on Mixed Model statistical analysis (see panel b), proteins were categorized into four groups: enriched by NTPs, enriched by transcription, reduced by NTPs, and unaffected "background" proteins. (**b**) Scatter plot shows $\log_2$ scaled +NTP/-NTP ratio of proteins bound downstream in the absence (x-axis) or presence (y-axis) of α-amanitin. Each circle represents a single protein, quantitated by averaging all peptides for that protein. Proteins are color-coded based on grouping. Proteins defined as "Enriched by transcription" (red) are to the right of the vertical dotted-line (95% confidence level for enrichment with NTPs) and below the diagonal blue dotted line (95% confidence level for inhibition by α –amanitin). Proteins "Enriched by NTPs" (blue) are defined as being over the 95% confidence level for NTP stimulation both with and without α –amanitin, but below the diagonal blue dotted line (95% confidence level for inhibition by α –amanitin). Proteins "Reduced by NTPs" (green) are over 95% confidence level for decreased binding in both conditions. Heatmaps show binding profiles for proteins 'Reduced by NTPs' (**c**) and 'Enriched by NTPs' (**d**). Relative reporter ion intensity for each protein was normalized to the untreated sample (-NTPs without $\alpha$-amanitin) set as 1.0. Proteins were then sorted and labeled by membership in a specific complex or functional group as listed to the right of the heatmaps. (e) Volcano plot of statistical significance (y-axis, -log10 of p-value) versus fold enrichment (x-axis, log2 of +NTP/-NTP ratio) for proteins associated with the downstream region of the transcription template in multiple replicates, as shown previously (Joo et al., 2017). A total of 712 proteins were identified in all three biological replicates each done with two technical replicates of quantitative mass spectrometry analyses on naked DNA templates. Each circle represents an individual protein. The TAFs, RNApII, Paf1C, and core elongation factors (Spt5, Spt6, and Spn1) are each highlighted as a different color. P-value is by t-test of specific protein values against total identified proteins. Dotted lines represent the 95% confidence level of changes.

Supplemental Figure 1

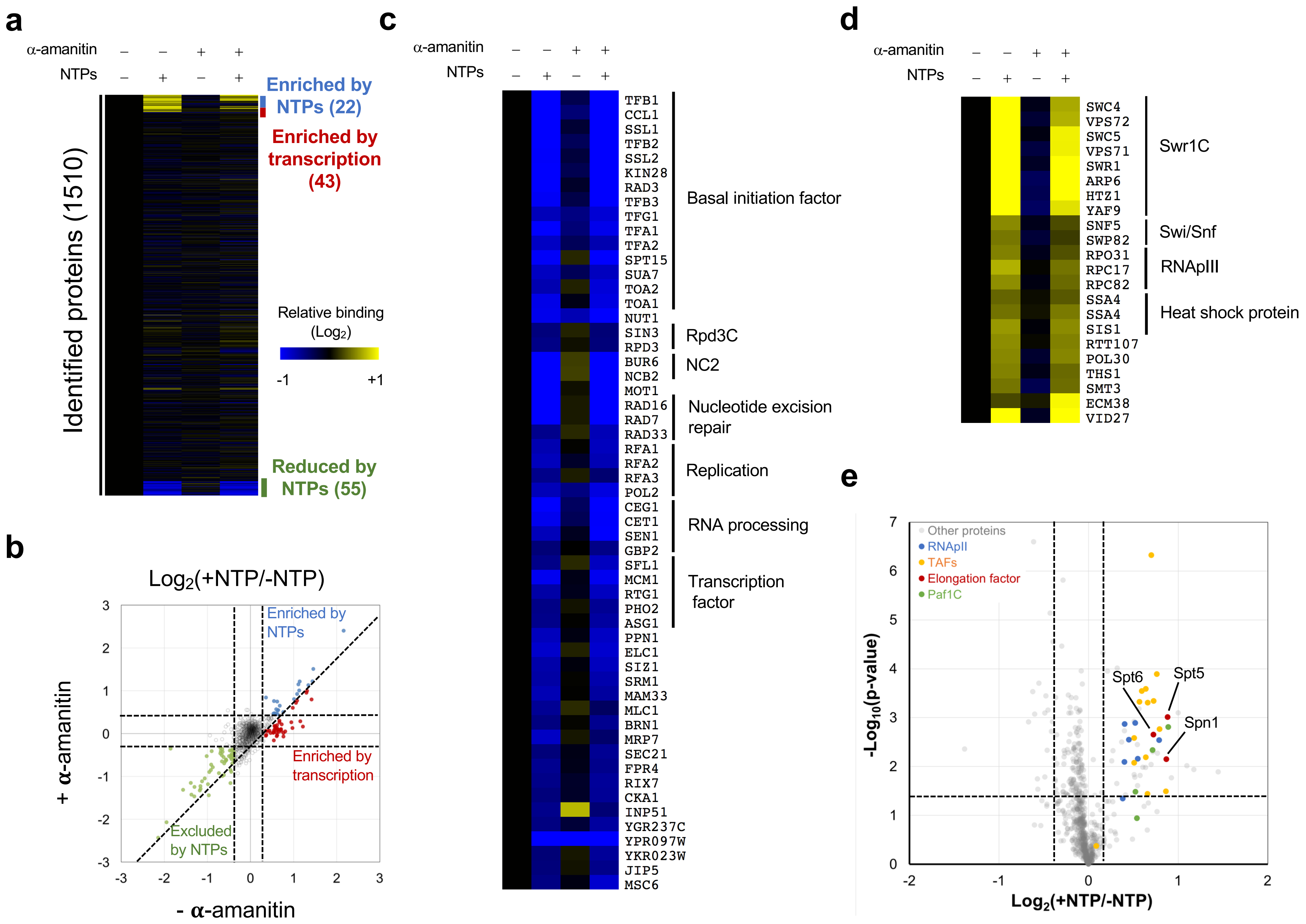

**Supplemental Figure 2,** related to Figure 2**.** Progression of CTD phosphorylation changes on stalled RNApII ECs.

(**a**) After PIC assembly with yeast nuclear extract and Gal4-VP16, transcription was initiated with NTPs (A, C, UTP and 3'-Ome-GTP). Downstream bound proteins were recovered by Ssp I digestion at 0, 1, 2, 4, 8, 16 min after transcription start. CTD phosphorylations were detected with specific antibodies against Ser5P (3E8) and Ser2P (3E10). Note that the lower Ser5P signal compared to **Fig 2b** is because that experiment used PstI to release both ECs and promoter-bound species.

(**b**) Different length G-less cassettes (+289, +128, and +90) were used for the time course CTD phosphorylation assay as in **Fig 2b**. After PIC assembly, transcription was initiated with NTPs (A, C, UTP and 3'-Ome-GTP). Total bound proteins were recovered at 0, 1, 4, and 8 min after transcription start. CTD phosphorylations were detected the same as in (**a**).

(**c**) After PIC assembly, NTPs (A, C, UTP and 3'-Ome-GTP; each 400 μM) or ATP (400 μM) were added in the reaction. Total bound proteins were recovered at 0, 1, and 4 min after the addition. CTD phosphorylations were detected as in (**a**).

### Supplemental Figure 2

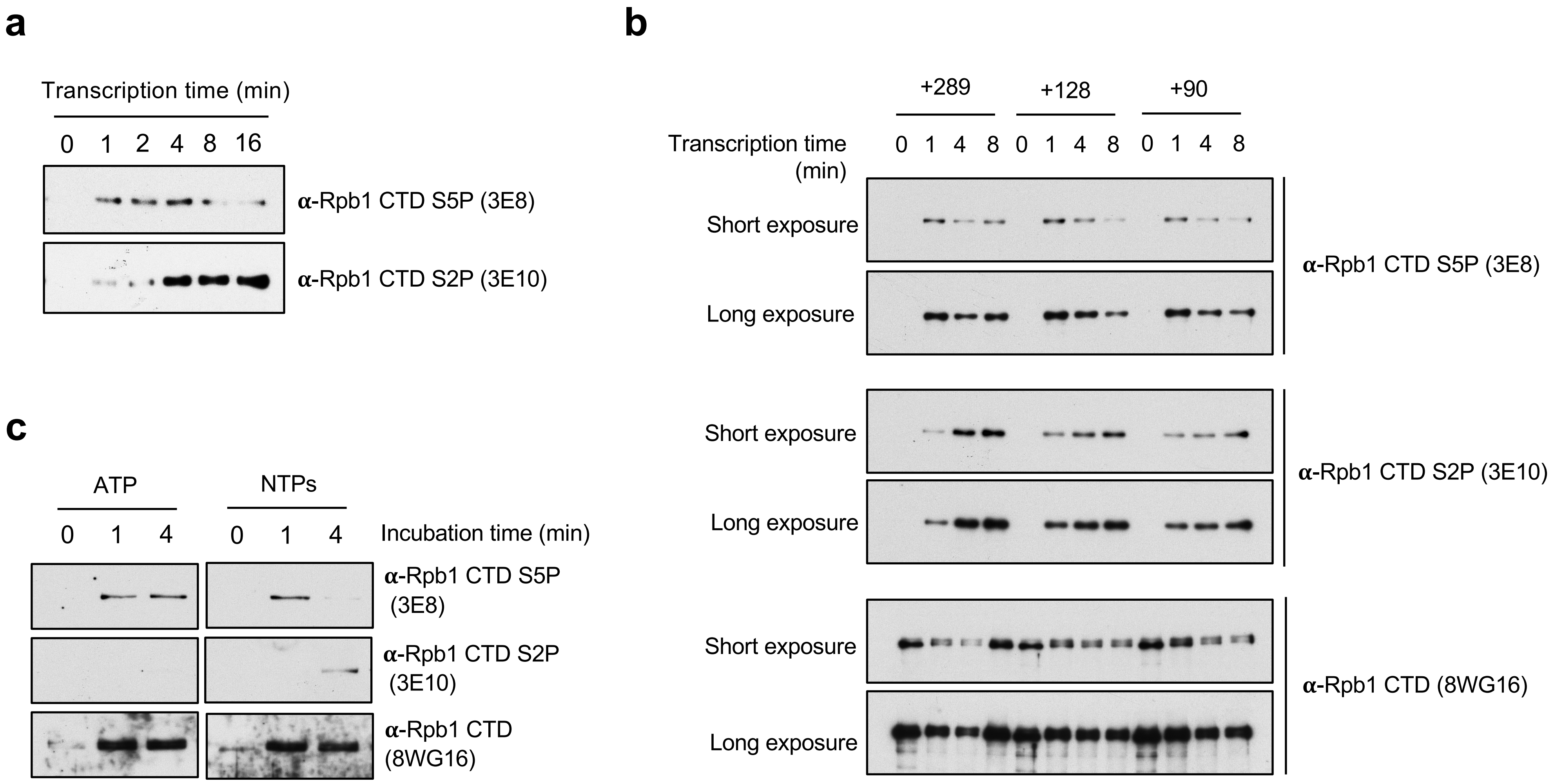

**Supplemental Figure 3**, related to Figures 3 and 4**.** Differential EC protein binding during transcription time course.

(**a**) Heatmap shows binding profiles for all identified proteins (total 1,098 proteins) from the quantitative mass spectrometry experiment shown in **Figure 3**. Relative reporter intensity of each sample was calculated, normalized to the untreated sample (0 min) as 1.0. Each line represents one protein. Proteins were clustered based on differential binding profiles into three groups (enriched by NTPs, reduced by NTPs, and unaffected). Early and late binding factors were identified by evaluating the slope of a line defined by the 1, 2, 4, 8, and 16 min time points. Green signifies an increase (more late binding) and pink an decrease (more early binding) over the time course.

(**b**) Mixed model plot of +NTP/-NTP ratio for each protein ($\log_2$ of averaged ratio at 1, 2, 4, 8, and 16 min). Dashed line shows cutoff for 95% confidence that protein levels are significantly changed by NTPs. The RNApII machinery (black) and core elongation factors (Spt4, Spt5, Spt6, Spn1, Elf1, and Paf1C subunits; blue) are color-coded.

(**c**) Heatmap shows binding profiles for proteins reduced by NTPs over time. Relative reporter intensity was calculated, normalizing to the background binding (0 min) as 1.0.

### Supplemental Figure 3

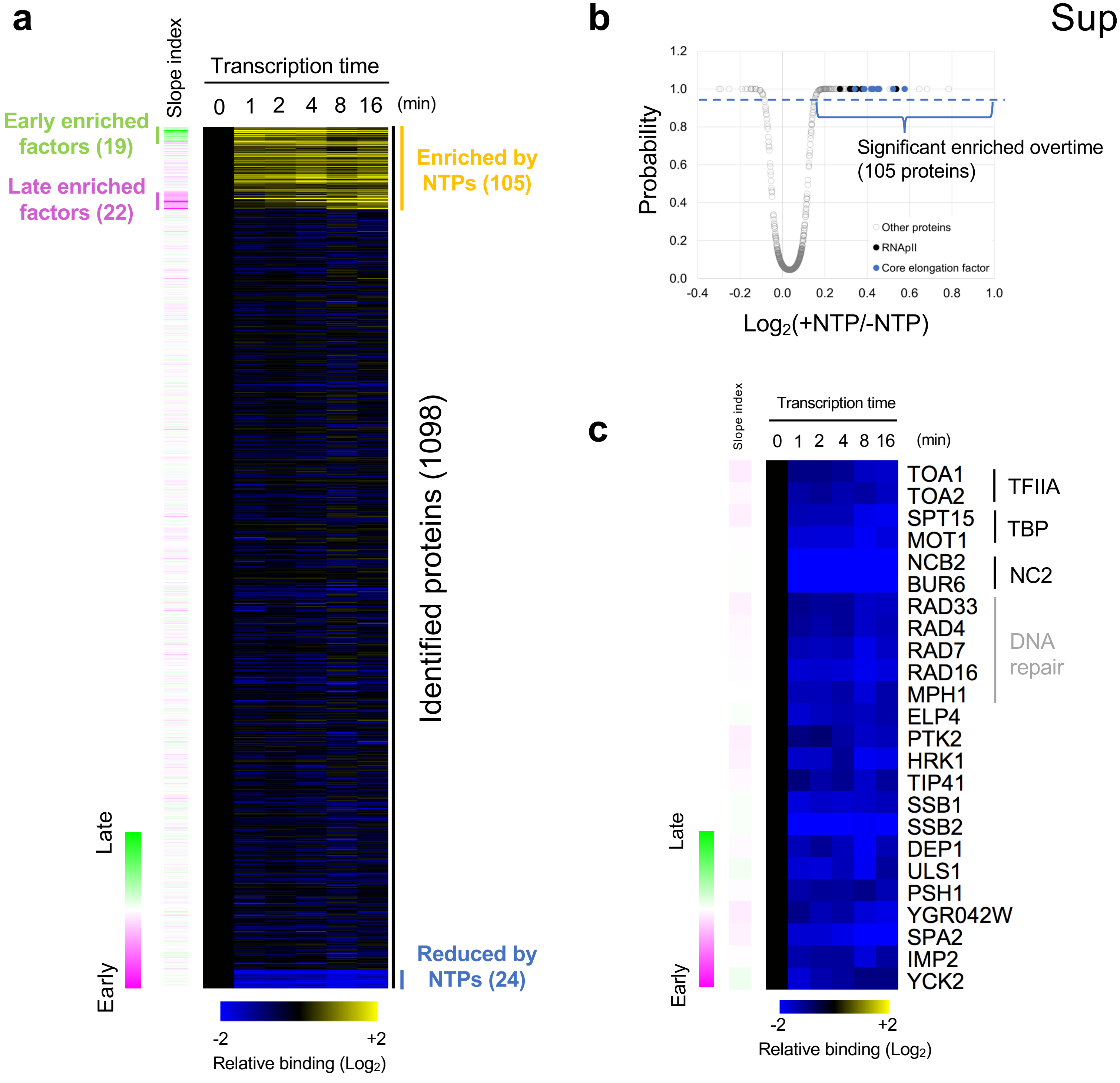

**Supplemental Figure 4**, related to Figure 5**.** Chemical inhibition of Kin28 kinase activity by CMK.

(**a**) Cell growth on plates with or without CMK inhibitor. The inhibitor sensitive *KIN28is* (YSB3356), but not wild-type *KIN28* strain (YF702), showed a severe growth defect in the presence of 1 μM CMK.

(**b**) Longer exposure panels are shown for Ser5P and Ser2P for the immunoblots from **Fig 5a**.

### Supplemental Figure 4

**a**

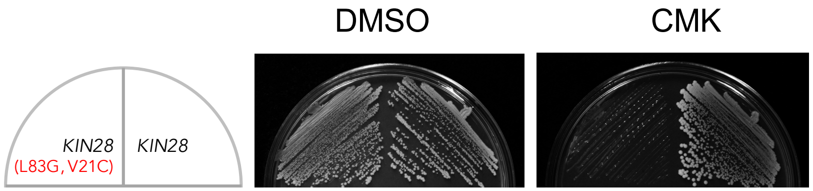


**b**

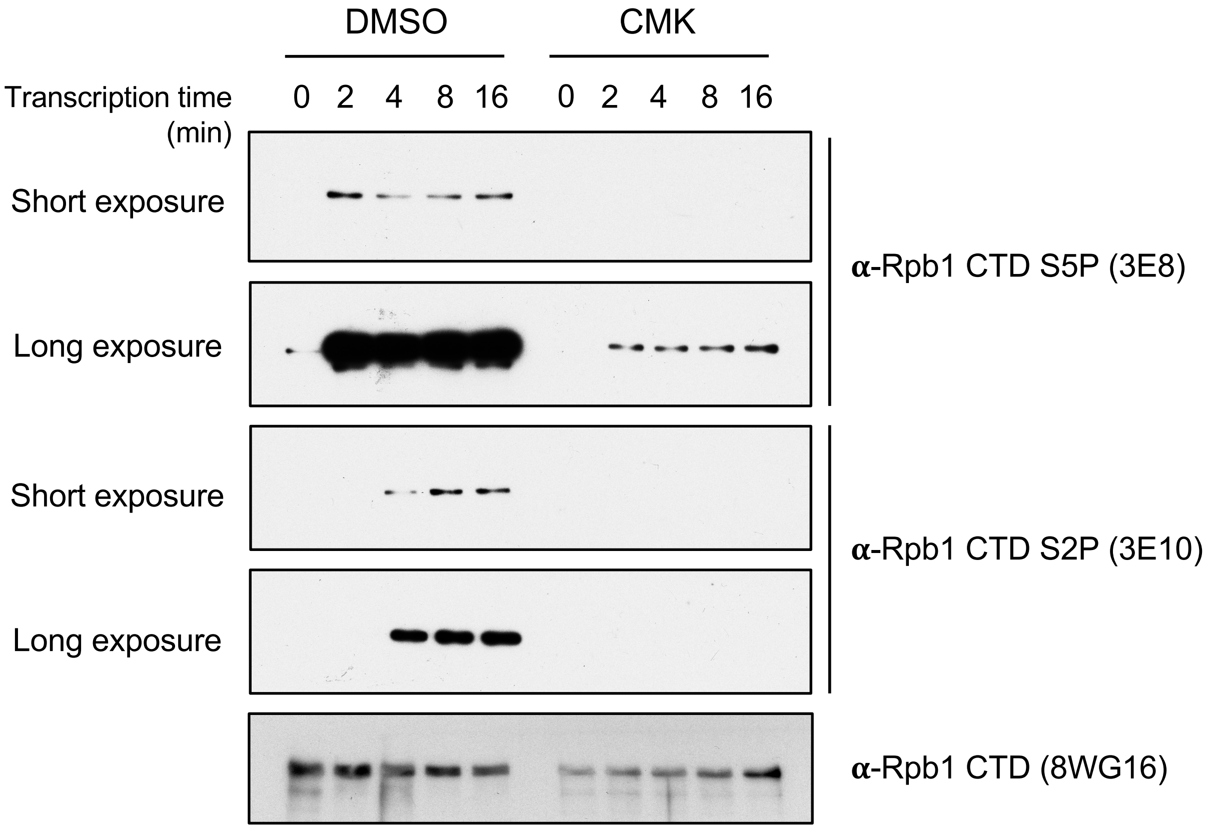

**Supplemental Figure 5**, related to Figure 6. Effects of Kin28 kinase inactivation on *in vitro* transcription.

(**a**) Control transcription reactions with nuclear extracts from the *KIN28* strain (YF702), showing no effect of CMK. After preincubation of extract with different CMK concentrations for 10 min, transcription was initiated by adding $^{32}P$-labeled NTPs for 30 min. Transcripts were recovered and detected by $^{32}P$ autoradiography on a 6% UREA-acrylamide gel. The level of transcription from proximal and distal start sites were measured by densitometry. Relative ratio of transcription was evaluated and graphed for each transcript.

(**b**) Differential effect of Kin28 inhibition on the CYC1 TSSs. *In vitro* transcription with nuclear extracts from the *KIN28is* strains (YSB3356) was performed in the presence or absence of CMK (500 nM) on two templates with different sized G-less cassettes:pUC18-G5CYC1 G- (SB649) and pG5CG-D2 (F916) (upper panel). Transcripts were recovered and detected by $^{32}P$ autoradiography on an 8M urea - 6% polyacrylamide gel. Ratio of relative transcription by CMK inhibition from proximal (#1 and #3) and distal (#2 and #4) start sites on each template were calculated using densitometry of three independent reactions. Standard deviation was evaluated with the student t-test and displayed in the graph as error bars. Note that the larger third band seen with each template represents read-through of the entire G-less cassette and therefore does not reflect initiation at the CYC1 promoter.

### Supplemental Figure 5

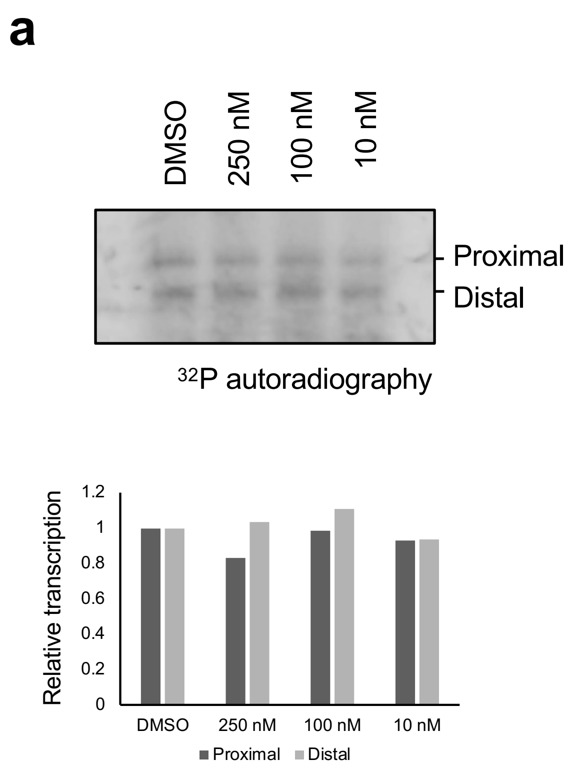


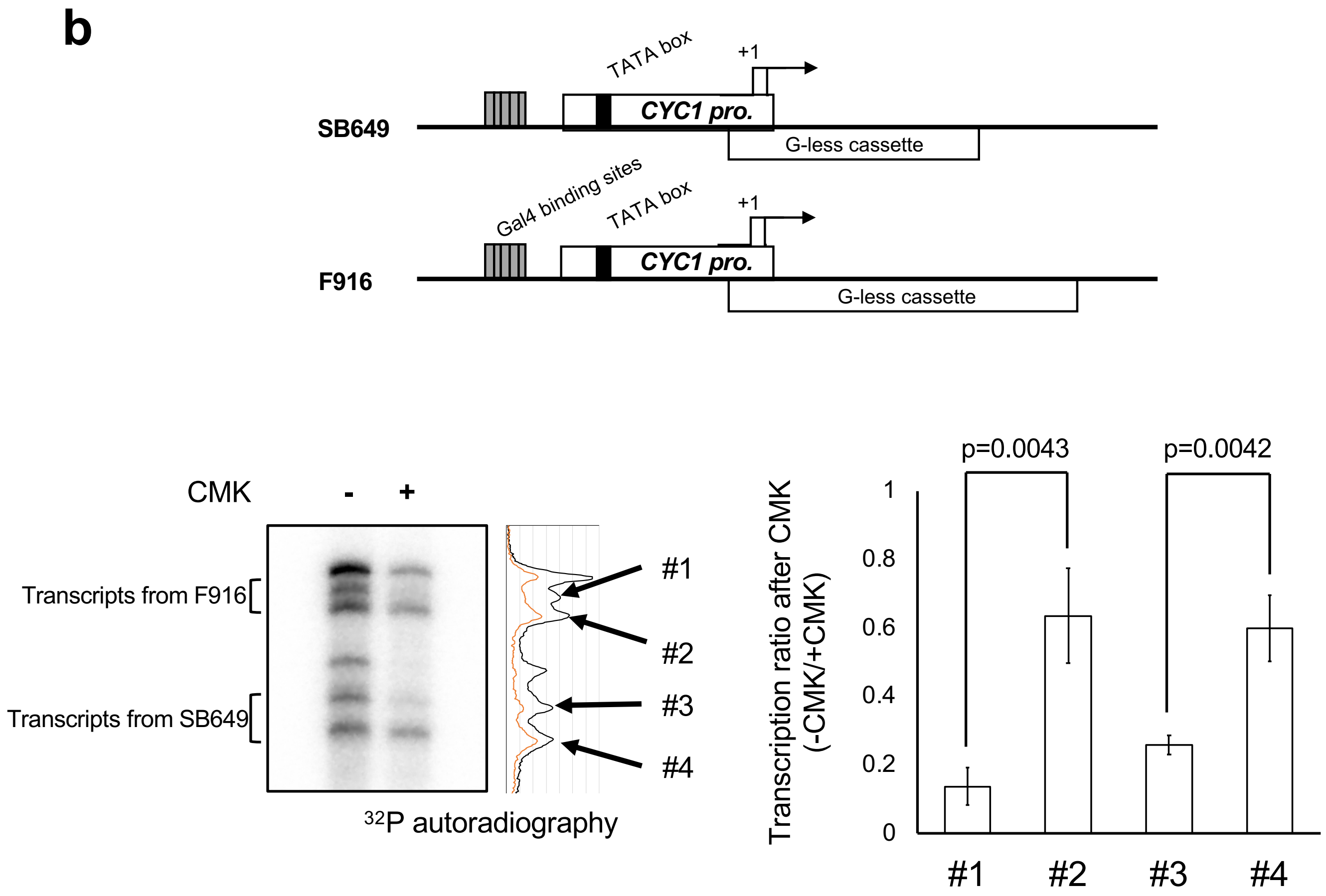
